# Overlapping perceptual benefits of temporal attention in aperiodic and periodic streams are mediated by delta phase alignment, but not alpha suppression, in sensory circuits

**DOI:** 10.64898/2026.09.11.750910

**Authors:** Christina Bruckmann, Assaf Breska

## Abstract

Predictions about the timing of upcoming stimuli allow to allocate attention proactively, facilitating perceptual processing. While both periodic and aperiodic predictions were associated with anticipatory suppression of occipital alpha-band activity, periodic predictions are thought to uniquely affect perception through entrainment of delta-band activity directly at the level of the sensory cortex. Here we isolated the perceptual effects of interval-based aperiodic predictions using a challenging perceptual paradigm with a delayed response. We compared the perceptual facilitation to entrainment-based predictions in isochronous streams, and how they are mediated by distinct neural mechanisms in delta and alpha bands. Compared to an irregular non-predictive condition, both interval-based and periodic temporal predictions facilitated objective discrimination performance and subjective visibility reports to an equal degree. Neurally, interval-based predictions led to increased delta-band phase alignment, equally to that observed in isochronous streams, with both effects centered on occipital regions. A computational model of oscillatory entrainment revealed that these results cannot be explained by entrainment to the aperiodic stream. Interestingly, anticipatory alpha-band suppression did not differ between predictive and the non-predictive conditions, dissociating the perceptual effects of temporal attention from modulations of occipital alpha amplitude. Overall, these results indicate that top-down temporal predictions can shape visual processing through phase alignment of non-oscillatory low-frequency activity at the sensory cortex, pointing to the context-independent neural mechanisms of temporal attention at the cortical level. More broadly, they highlight the unique functional and neural features of temporal attention.

## Introduction

Our senses are continuously confronted with a dense flow of incoming information. To allocate resources optimally, the brain prioritizes moments in time based on temporal regularities in the environment. Regularities such as isochronous rhythms, repeating sequences, and cue-interval associations have been shown to guide attention, resulting in facilitation of both motor performance and perceptual processing at the attended moments (e.g., intervals: (1,2); rhythms: (3,4); sequences: (5–7)). However, whether and how distinct neural mechanisms mediate temporal prediction and the subsequent attentional modulation of perceptual sensitivity in different temporal contexts remains controversial.

Traditionally, temporal prediction in aperiodic streams was explained by associative learning of a consistent interval between two events (cue-target). Initiated by the cue, time is tracked by centralized interval-timing mechanisms, e.g. a pacemaker-accumulator (8), and attention is increased top-down to peak when the interval elapses. In contrast, in (quasi)rhythmic streams, as present in music or biological motion, entrainment models suggest a more bottom-up mechanism, where attention is guided in time by aligning low-frequency neural oscillations of the sensory cortices with the stream of rhythmic input (9,10). This way, the optimal oscillatory phase for perception coincides with upcoming stimuli. Findings of ramping neural activity in aperiodic streams (11,12), thought to reflect pacemaker-related accumulation or the gradual increase of attention, and findings of neural phase alignment in periodic streams (13), were taken as evidence of this mechanistic dissociation. However, recent work (11,14) revealed that in a speeded reaction time task, cued-interval predictions in aperiodic streams lead to similar reaction time benefits and adjustments of ramping activity as in rhythms. Strikingly, phase alignment to the optimal angle at fronto-central circuits reached a similar magnitude in both contexts. This overlap in phase alignment patterns challenges the traditional taxonomy, and calls into question the assumed role of oscillatory entrainment in rhythmic predictions.

However, how these neurocomputational principles extend to facilitation of perceptual processing by temporal predictions remains an open question. The use of a perceptually non-demanding motor task in previous work may have discounted behavioral and sensory processes that are only engaged under high perceptual load, while emphasizing the role of frontal motor circuits. First, at the sensory processing level, rhythmic streams directly elicit phase entrainment at input-specific sensory circuits (13), potentially providing perceptual benefits for rhythm-based relative to interval-based prediction. However, it is not known whether interval-based prediction can drive phase alignment in sensory circuits. Conversely, rhythmic and aperiodic prediction lead to similar anticipatory alpha suppression in occipital areas (11), a well-established marker of visual attention (15,16), suggesting that sensory circuits are modulated across temporal contexts. Second, at the level of fronto-central circuits, it is not clear whether previously observed phase alignment in frontal circuits is specific to motor preparation, or reflects task-independent involvement of control networks. It has been suggested that in interval-based predictions this could reflect temporally consistent ramping activity (11), while in rhythm predictions, motor-based time representations (17). Isolating the architecture of temporal prediction in purely perceptual settings is essential for dissociating rhythm- and interval-based prediction, but also more broadly, for understanding attention and how top-down and bottom-up processes converge in sensory circuits.

Another important gap in our understanding of temporal attention, especially in the context of perception, is how it impacts different aspects of perception. The majority of studies reporting perceptual benefits of temporal attention quantified perception through objective measures, such as discrimination performance, with few studies investigating the influence temporal attention has on subjective perception (e.g. (18). This differentiation, while central to consciousness research, has been largely overlooked in the timing literature. To our knowledge, only two studies directly assessed the impact of temporal attention on both of these facets of perception (2,19). In an interval context, these studies found a benefit of temporal attention on both measures, with Ciupińska and colleagues also reporting an interaction between objective and subjective measures in their task that included both spatial and temporal attention. Yet, given that neural entrainment is hypothesized to drive attention exogenously in addition to top-down memory effects, the putatively different neural mechanisms involved in rhythmic timing might affect these aspects of perception differently.

The current study combined psychophysics, electroencephalography (EEG) and computational modelling to investigate the context-specific mechanisms of temporal prediction, and their attentional impact on multiple aspects of perception. Specifically, we aimed to uncover how top-down interval-based predictions impact low-level sensory processing, and compare these purely top-down generated predictions with rhythmic predictions, which directly act on the sensory cortex. Accordingly, we designed a demanding visual discrimination task in which a visual target was preceded by cue stimuli with either a rhythmic or an interval structure, providing strong temporal predictability, or by irregularly timed stimuli as control. Critically, to minimize motor preparation in the target anticipation window, perceptual judgments were only reported after a significant delay, rather than in speeded reaction to target onset. To obtain a full picture of perceptual effects of temporal attention, participants reported both the orientation as well as the subjective visibility of the target, for a wide range of target contrasts that ensured sensitivity to modulations of both objective performance and subjective visibility. Finally, we used a computational time-resolved model of entrainment to test the predictions of this mechanism in these different temporal contexts.

## Materials and Methods

### Participants

Thirty-nine healthy volunteers with normal or corrected-to-normal vision and no history of neurological disease were recruited to participate in two experimental sessions - one behavioral, one with EEG - on two separate days. All participants provided written informed consent and received monetary compensation. Eight participants were excluded from analysis as they did not return for the second session. One participant was removed from behavioral analysis only due to errors in data acquisition in the first session. Two more were rejected from all analyses due to predefined rejection criteria as outlined below. The final sample thus includes 29 participants for the EEG data (2 left-handed; 19 females, mean age: 23.7 years (SD: 2.27)) and 28 for the behavioral data. The study design was approved by the ethics committee of the medical faculty of the University of Tübingen, in accordance with the Declaration of Helsinki.

### Stimuli and Design

#### Set-Up and Stimuli

All stimuli were created with custom code in MATLAB R2024b (The MathWorks Inc., Natick, Massachusetts) and displayed using Psychtoolbox-3 (20) on a CRT monitor with a resolution of 1024×768 pixels, a pixel density of 65.02 dots per inch and a refresh rate of 60Hz. Responses were given through keyboard button presses and a chin-rest was used to fix the viewing distance at 90cm.

Participants performed a perceptual discrimination task (Fig. 1A), reporting the orientation (left/right) and subjective visibility of a visual target. The target was a grayscale Gabor grating (spatial frequency: 2.8 cycles/degree; Gaussian envelope standard deviation: 3.0°), tilted by 45 degrees either to the left or the right from the meridian, and presented for 16.6ms at one of 10 different contrast intensities (individually selected for each participant, see below).

**Figure 1.**
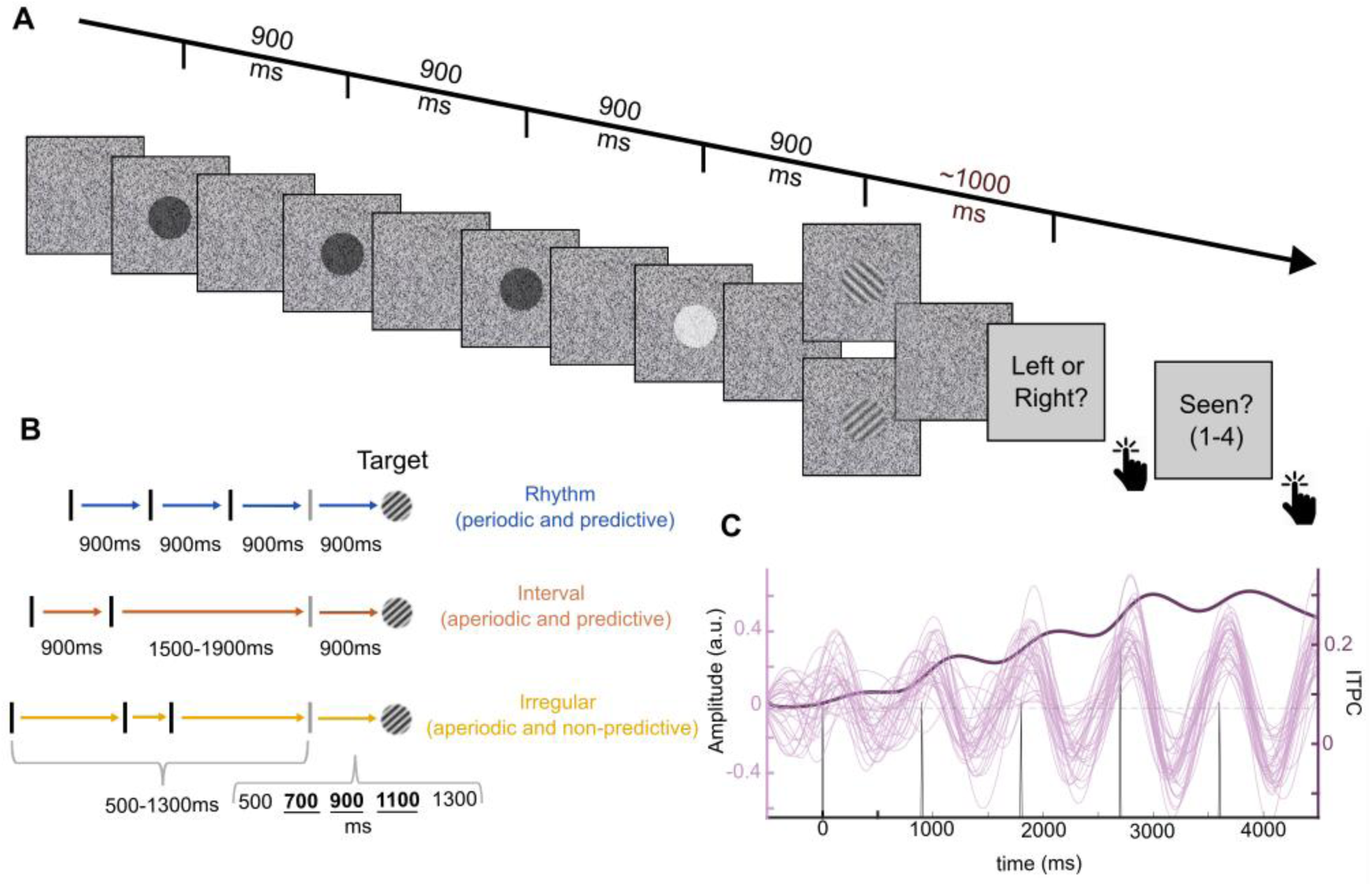
Experimental paradigm and oscillatory entrainment model. **A.** Illustration of a Rhythm trial. Continuous dynamic visual noise mask, with superimposed cue and warning signal stimuli. Target was a grating tilted to the left or to the right, embedded in the noise mask with varying intensity. Delayed response paradigm. All stimuli were presented in the center of the screen. **B.** Experimental conditions.

Each target was preceded by black circular cue stimuli (diameter: 120px ≈ 4.7cm, viewing angle: 3°, duration: 100ms) and a white circular warning signal (WS, same size and duration), indicating that the next stimulus would be the target. The entire stimulus sequence was embedded in a continuous dynamic noise mask (300 x 300px ≈ 11.72 x 11.72cm, viewing angle: 7.5°), presented throughout the duration of the trial. The mask was created by setting a random luminance value (0-100%, 255 steps, 8-bit monitor) for each pixel and changing it every monitor refresh cycle (16.6ms). While the cue stimuli were superimposed over the mask, rendering them clearly visible, the target stimulus was blended into the mask, with the visibility changing based on the desired target contrast. The mask presentation started 1000-1600ms (randomly chosen, uniform distribution) before the first black cue stimulus and ended 1000ms after target offset. To reduce interference of motor preparation during the anticipation stage, participants did not provide a speeded response following target onset. Instead, the prompt to report the orientation of the target stimulus appeared 850-1250ms after mask offset, and any responses were registered only after each question was presented, thus at least 2s after target onset. Participants were required to provide an orientation judgment even when subjective visibility was low. As soon as participants responded to the orientation question, the question regarding target visibility appeared on screen. Subjective visibility was recorded with the four-point Perceptual Awareness Scale (21,22), using the four visibility levels of “no experience”, “brief glimpse”, “almost clear experience”, “clear experience”. The PAS was chosen because its lowest visibility level has been shown to closely align with chance performance (23). Participants had no limit on their response time but were encouraged to not overthink their answers. After three seconds of no response, a message was presented urging the participants to respond.

#### Experimental conditions

The main experiment consisted of three conditions, presented in separate blocks (Fig. 1 B). Each trial of the Rhythm condition consisted of 3 black cue stimuli, WS and target, all with a stimulus onset asynchrony (SOA) of 900ms, making target timing predictable by synchronizing to the emergent beat (1.11 Hz). In the Interval condition, a pair of black cue stimuli appeared with a 900ms SOA, followed by the WS and target, also with a 900ms SOA, making the target timing predictable based on memory of the cue interval. The SOA between the second cue and the WS was randomly jittered between 1500ms and 1900ms, approximating the timing of the rhythm condition while strongly reducing the capacity of an oscillator to entrain to the sequence (for details see “Computational Modelling” below). Finally, as a control condition with reduced temporal predictability, the Irregular condition consisted of 3 black stimuli, WS and target. The SOAs between consecutive cue stimuli, as well as between the last cue stimulus and the WS, were randomly chosen from a uniform distribution between 500ms and 1300ms (50ms steps), with the constraint that consecutive ISIs within one trial had to have at least a difference of 200ms. The SOA between WS and target was varied across trials but had a mean duration of 900ms, matching the predictable target interval in the other conditions. An equal number of trials was presented with a WS-target SOA of 500ms, 700ms, 900ms, 1100ms and 1300ms.

Rhythm (blue) had a fixed 900ms SOA, Interval (red) repeated a 900ms interval with jittered inter-interval SOA, Irregular (yellow) had a high variability in the interval SOA. Bold text: target intervals that were analyzed in comparison to the predictive conditions. **C.** Simulations of the oscillatory entrainment model. Pink: individual trial dynamics in the Rhythm condition; Purple: inter-trial phase concentration (ITPC). An entrainment-based process leads to gradual increase in ITPC as the stream progresses.

To estimate the full psychometric curve, targets were presented with 10 different contrast intensities, with 12 trials per intensity level, resulting in 240 trials per condition across the two sessions. In the irregular condition, the 240 trials were split evenly across the central 3 WS-target SOAs (700, 900, 1100), resulting in 80 trials across sessions for each SOA. In addition, to balance the contrast values and the number of trials at each possible SOA, 160 additional trials (10 contrasts x 8 trials x 2 sessions) were presented across the two outermost SOAs (500ms and 1300ms) resulting in a total of 400 trials. This was done to ensure that the contrast levels and trial numbers are equivalent to the predictive conditions in subsequent analyses (see below). In the second session, 36 ‘catch’ trials without target presentation were added to each of the two predictive conditions and 40 were added to the irregular condition. No questions regarding target perception were presented for these trials.

### Procedure

The experiment was conducted in an electrically shielded, dimly lit, and sound attenuated chamber, in two sessions across two separate days (days between sessions: 3-55 days, mean 12.8). The conditions were presented in a block-wise manner (3 blocks for each predictive condition, 4 blocks of irregular), To minimize order effects while respecting the unequal condition frequencies, the first block was always of the Irregular condition, followed by three cycles of Rhythm-Interval-Irregular, or Interval-Rhythm-Irregular (counterbalanced across participants).

#### Training

The first session began with task instructions and several (∼3-10) preliminary practice trials, to familiarize the participants with the discrimination task and the perceptual awareness scale. All trials given at this stage only included the continuous noise mask with embedded targets, with no temporal cues or WS. During this stage, participants were trained on the PAS according to its guidelines (22,24). After confirming that participants understood the different PAS categories, they completed a staircasing procedure to adjust target intensities accordingly (see below).

Next, participants performed at least 5 practice trials, this time from the irregular condition, to familiarize themselves with the cues and warning signal. The target in the first trial was always shown at full intensity, while the target contrast for the other trials was chosen randomly from the values determined by the staircase.

The first Rhythm and the first Interval block were preceded by an explanation of the respective temporal structure. Participants were encouraged to actively utilize the temporal cues to predict the moment of target onset. To ensure that all participants could explicitly identify the temporal regularity, they performed a 2-alternative-forced-choice task before the first block of each predictive condition. In this task, participants were presented with two consecutive streams, in which the cue matched the respective predictive condition, and the target was vertical and presented in maximum visibility. The target appeared at the cue-predicted time in one of the streams, and too early or too late in the other stream. The task was to report in which of the two streams the target appeared at the cue-predicted moment and ended when participants answered correctly four times consecutively. In case of errors, further clarification was provided.

In the second session, participants only received a short explanation to remind them of the temporal structures, but did not complete any practice trials without cues. Instead, they performed the staircase procedure right away, before completing 5 or more practice trials of the irregular condition as a reminder. The second session did not include the 2AFC task before the predictive conditions.

#### Staircase Procedure

Even though we used a range of 10 target contrasts, we still opted to use an adaptive procedure, to ensure coverage of both the subjective and the objective psychometric function without temporal expectation, as well as allow us to detect their potential shifts when attention is temporally guided. As such, precisely estimating a specific threshold was not necessary, but instead we approximated the 50% threshold of subjective visibility to center the experimental contrasts around that value. To this end, in each trial participants viewed a target embedded in a continuous noise mask (no cues or WS), and reported both its orientation and their subjective visibility of the target as in the main experiment. Contrast levels of the target were adjusted after each trial using a one-up-one-down staircase procedure. The response evaluation combined the objective and subjective tasks, such that a response was considered positive if orientation judgment was correct together with a PAS rating higher than 1 (1= “no experience”), and negative otherwise. The staircase terminated automatically after 40 reversals. The subjective 50% threshold was estimated as the mean contrast level of the last 10 reversals.

This led to the 50% subjective perception level approximately coinciding with contrast level 5. This way, we assured that the contrast range included strong enough values to sustain the motivation of the participant and their belief in the predictability of the cues, and weak enough values to probe facilitation by temporal attention.

#### Post-Study Questionnaire

After each session, participants responded to a short questionnaire. This questionnaire contained two questions that explored the strategies they had used during the task (one open, one with multiple options, data not analyzed). Lastly, participants were asked on a scale of 0-100 how well they perceived that they had managed to use the rhythm and interval cues. To avoid report bias, we assured the participants that the responses to these questions were anonymous and would not affect their payout.

### Behavioral Analysis

Behavioral data were analyzed in MATLAB R2024b (The MathWorks Inc., Natick, Massachusetts) using custom scripts, the statistics toolbox and the *psignifit 4* toolbox (v2.5.6, (25). For each participant, performance and visibility data were aggregated across the two sessions. Additionally, for the irregular condition, only trials with target intervals of 700ms, 900ms, and 1100ms were analyzed, making the remaining trials more comparable to the predictive conditions and matching the total number of trials across conditions. In line with previous studies (2) the PAS ratings were binarized, with 1 being scored as ‘not-perceived’ and anything above being considered as ‘perceived’ (for non-binarized analysis, see Relative Psychometric Function Analysis). Subsequently, for each participant psychometric functions were fitted separately for objective performance and subjective visibility within each condition, across contrast levels of the target. We fitted a four-parameter logistic function (as implemented in psignifit4) to the subjective visibility ratings, allowing threshold, slope, and lapse rate to vary freely. For the objective task, the guess rate was fixed at 50%. For visualization purposes, we also computed group-level psychometric curves by averaging the proportion of correct responses at each stimulus level across participants and fitting a curve to these aggregated data. All statistical analyses were conducted using the resulting parameter estimates from the psychometric functions of the individual participants.

#### Rejection Criteria

We excluded participants violating the following criteria set beforehand to ensure data quality: not using all responses of the subjective awareness scale (N=0) or having an irregular threshold below contrast 2 or above contrast 8, indicating incorrect staircasing (N=1). Statistical outliers were identified using an interquartile range (IQR) criterion, excluding participants whose threshold shifts between conditions exceeded ±2×IQR from the first or third quartile (N=1). Thus, 27 participants were included in the following analyses.

#### Statistical Analysis

To determine whether temporal expectation impacts the perceptual threshold, psychometric thresholds (75% correct for objective performance and 50% ‘seen’ for subjective visibility) were extracted for each participant, condition, and perceptual measure. A repeated measures ANOVA was conducted to assess the effect of condition on perceptual threshold, separately for subjective and objective measures. Subsequently, planned contrasts of thresholds were evaluated using one-tailed paired samples t-tests, assessing the hypotheses that temporally informative cues (Rhythm and Interval conditions) lead to a lower threshold than less predictive ones (Irregular), and that rhythmic cues would lead to the largest shift due to low level entrainment. To examine whether temporal expectation has an influence on the slope of the psychometric function, slope values were extracted at the psychometric threshold for subjective visibility (50%) and objective performance (75%) respectively. This was done for each participant and condition separately. A repeated measures ANOVA was done to assess the effect of condition on the slope of the psychometric function. As measure of effect size across all analyses, we report Cohen’s *d* for t-tests and *η²ₚ* for ANOVAs. We also report Bayes Factor (*BF*) to quantify evidence against hypothesized directional effects (e.g. superiority of rhythm-based prediction).

#### Relative Psychometric Function

To assess the relation between subjective visibility and objective performance under different attention conditions, we used the relative psychometric function and the associated toolbox developed by (26). For this, the objective performance curves were fitted using the Weibull function with a fixed guess rate at 50%, and a fixed lapse rate at 0.02, using a maximum-likelihood estimation. We estimated the subjective functions using the PAS scores binarized as above, fitting a Weibull function using a maximum-likelihood estimation without fixed parameters. Participants for whom any of the six functions resulted in infinite slopes (N=5) were rejected from further analysis. We subsequently fit the relative psychometric function by fitting the subjective visibility as a function of objective performance and extracted the area under curve (AUC) for each participant and each condition. The AUC was compared between conditions using a two-tailed repeated measures t-test.

To increase comparability to recent results from spatial attention (27), we repeated the analysis using the entire, non-binarized, PAS scale as mean ratings and fitted a scaled Weibull function using least-squares minimization of the sum of squared errors, without fixed parameters for each subject and condition. With the graded visibility scale, two participants for whom any of the six functions resulted in infinite slopes were rejected from this analysis.

### EEG Recording and Preprocessing

Electrophysiological data was recorded continuously with an ActiveTwo system (Ag/AgCl pre-amplified electrodes, BioSemi, The Netherlands), using 64 scalp electrodes mounted on an elastic cap according to the extended 10-20 system. Further electrodes were placed near the tip of the nose, above and below the right eye, bilaterally near the lateral canthi of the eyes, and on the mastoid bones. The signal was sampled at a frequency of 1024 Hz (24 bits/channel), with an 5th order anti-aliasing low-pass filter (204 Hz).

The neurophysiological data were imported to MATLAB R2024b (The MathWorks Inc., Natick, Massachusetts) preprocessed with custom scripts Excessively noisy channels were identified through visual inspection or excessive artifact (see below) and later interpolated based on the mean activity of the nearest neighbor electrodes. The raw data was detrended using a smoothed end point, re-referenced to the nose and filtered with a 0.1 Hz butterworth high-pass filter as well as a notch-filter at 50hz to remove line noise. Ocular artifacts were identified through independent component analysis (ICA) and removed based on visual inspection of the scalp topography and component time course. Epochs containing non-ocular artifacts were automatically marked using the rdi-function from EEGLab on 50Hz low-pass filtered data based on the following criteria: change of amplitude more than 100μV within a 200ms window, less than 0.5 µV amplitude change within 200ms, as well as activity exceeding + or - 100μV. The automatically selected artifacts were subsequently evaluated through visual inspection.

### EEG Analysis

EEG data were analyzed in MATLAB R2024b (The MathWorks Inc., Natick, Massachusetts) with custom scripts and the CircStat toolbox (28).

#### Delta Inter-trial phase Concentration

Delta-band inter-trial phase concentration (ITPC) was calculated separately for an occipital electrode cluster (Oz, O1, O2, POz, PO3, PO4, PO7, PO8) with nose-referenced data, and a central electrode cluster (FCz, FC1, FC2, Cz, C1, C2) using the average mastoid signal as a reference. For each, delta-band phase was extracted from the entire time series by band-pass filtering the data (Butterworth, 1.11 Hz ± 0.75 octaves, 24 dB/octave roll-off). The Hilbert transform was applied to the filtered signal using MATLAB’s built-in function, and the instantaneous phase was obtained from the resulting analytic signal.

Subsequently, the data were segmented from −700ms to 1200ms around the warning signal separately for each condition and averaged across electrodes. For the irregular condition, only trials with targets appearing at 900ms or later were analyzed to exclude interference of target-evoked activity in the window of interest. The data was averaged across trials and baseline-corrected to the expected chance-ITPC which was obtained by generating random angles for each trial of each condition, averaging them, and repeating this procedure 10,000 times per condition.

To compare the average pre-target IPTC across conditions, we conducted directional within-subject t-tests on the activity 800-900ms post WS, hypothesizing that temporal prediction would lead to increased IPTC, and rhythm more so than interval. This was done separately for the occipital and the central cluster. Further, we conducted cluster-based permutation analyses (29); 10,000 permutations) on the same contrasts from 0ms to 900ms post warning signal. Lastly, verified that the results were not explained by those irregular trials in which the target appeared later than 900ms and would thus not lead to target-evoked responses around the 900ms window, potentially lowering the IPTC in the window of interest.

#### Computational modelling

To obtain predictions of delta-band dynamics in the three experimental conditions based on an oscillatory entrainment mechanism, we simulated this process using a time-resolved coupled-oscillator dynamical system model (Fig. 1C). Based on (11), we focused on an oscillator whose frequency matched the mean rhythm frequency of 1.11 Hz, which also matched the predictive interval. The dynamical system included two state variables, the instantaneous phase angle (*θ*) and amplitude (*r*):

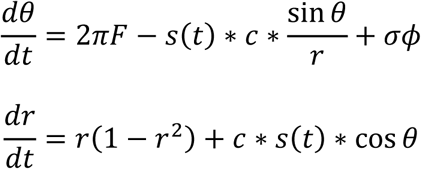

In this formulation, *F* denotes the natural frequency of the oscillator, *s*(*t*) denotes the instantaneous energy level of the external stimulus at time *t*, *c* denotes the coupling strength between the oscillator and the external input, and *σ* denotes the standard deviation of gaussian angle noise *ϕ*, therefore determining the rate of dephasing.

In the absence of external input (s=0), this results in self-sustaining limit-cycle dynamics, oscillating in frequency *F* and stabilizing at *r*=1. When an external stimulus is introduced (*s*>0), the system is phase-reset towards an angle of *θ*=0, defined here as the optimal phase, with the strength proportional to *c*, resulting in a momentarily increased phase concentration across iterations irrespective of the pre-input phase, before, in the absence of further external stimulation the system dephases at the rate indicated by the dephasing parameter. To simulate this model for a specific [*θ*, *r*] starting point and given set of parameters, we numerically integrated the dynamical system using a fourth order Runge-Kutta algorithm with a fixed integration step (MATLAB function ode4.m), to avoid missing time points in which stimulus energy is different than 0.

To model our experimental paradigm, we recreated the stimulus streams of the different conditions by introducing external stimulation to the system. This was implemented as a time resolved vector with a sampling rate of 100 Hz (10ms steps), that contained a non-zero value at the time points in which stimuli were presented in the experimental streams, and zero otherwise. To account for approximate conduction delays from stimulus presentation on screen to visual perception, the non-zero values were set not at stimulus onset but with an additional delay of 50ms. This time parameter was set to align the ITPC peak between the model EEG data, and changing it did not affect the pattern of differences between conditions. To account for the increased stimulus energy of the white warning signal compared to the black cue stimuli, we set the stimulation strength as 1 for the WS and 0.7 for cue stimuli. The simulated data was created by running 29 simulated participants, matching our participants sample size, each consisting of 150 trials within condition, matching the trial count of our paradigm. In each trial, the oscillatory system was initiated at a random phase and subsequently exposed to the stimulus stream, and the resulting radius and angle values were combined to create a simulated EEG signal as: *EEG* = *r* · cos(*θ*). We then bandpass-filtered the resulting simulated data around the natural frequency, as was done for EEG data, and calculated the ITPC across the 150 trials for all time points from 700ms before the warning signal to 300ms post-target time, resulting in one ITPC time course for each iteration and condition. Lastly, the ITPC was averaged across iterations and corrected for the ITPC expected by chance, estimated by calculating the ITPC of 150 randomly chosen angles, to match our trial count (repeated and averaged across 1000 times for stability).

As system dynamics, and potentially condition differences, could depend on the entrainment strength and dephasing rate parameter values, we took two approaches. First, we fit these two parameters to the experimental results obtained for the Rhythm condition (c=0.5, σ=2.8) such that the delta-band ITPC of the model would match that observed in the Rhythm condition in both the pre-warning signal and pre-target stimulus windows. We subsequently used the same parameters on a stimulus stream matching the interval condition, to estimate the expected stimulus-driven phase alignment in the interval condition. Second, we performed an exhaustive grid search of the parameter space to examine whether any possible combination of parameters could account for the observed phase-alignment in both the Interval and the Rhythm condition purely through an entrainment-based mechanism. For this, we ran the entire simulation protocol multiple times, cycling through the parameter space (in a range that leads to ITPC strength in the range of the values observed in the data and typically in the literature,(14,30,31)) of both entrainment strength (c=0.1:0.1:1) and dephasing rate (σ=1:0.2:3). We then compared the resulting simulated difference in delta-band ITPC between the Rhythm and Interval condition to the EEG data results, asking whether some parameter combination leads to stronger ITPC for rhythm and baseline level for interval in the 100ms pre-WS window, and a stronger than baseline but matched across conditions ITPC in the 100ms pre-target window.

#### Event related potentials

The event-related potentials analysis focused on ramping of negative potential in a frontocentral cluster (FCz, FC1, FC2, Cz, C1, C2), where it was previously observed in temporally predictive contexts (often referred to as Contingent Negative Variation, CNV, (11,12,32)). As our goal was to examine the co-occurrence of ramping activity and ITPC effects which we expected to find also in occipital regions, this analysis was also conducted in the occipital cluster used for the ITPC analysis (Oz, O1, O2, POz, PO3, PO4, PO7, PO8). The broadband data were segmented from 200ms before to 1100ms after the warning signal, and averaged across artifact-free trials. To test the existence of negative ramping activity, we first compared the amplitude averaged across a time window of 100ms pre-target to the baseline time window (averaged −200 to 0 before the WS), using one-tailed t-tests. Where negative ramping from baseline was found, we tested for condition differences in the pre-target window using paired t-tests. To examine potential differences in ramping slope, we fitted a linear model to the data from 600ms to 900ms post-WS of each condition, compared the slopes to 0 using one-tailed t-tests, and between conditions using paired t-tests. Rather than fitting to individual participant data, to obtain the slopes we used a jackknifing procedure, with appropriate corrections to statistical tests (33).

#### Alpha Amplitude

Pre-target alpha-band amplitude was analyzed for an occipital region of interest (Oz, O1, O2, POz, PO3, PO4, PO7, PO8). For the irregular condition, only trials with targets appearing at or after 900ms were included to exclude trials with interference of target evoked potentials in the window of interest, while still matching the number of trials across conditions. The data were segmented from 400ms before to 1500ms after the warning signal and padded with 500ms at each end to absorb edge artifacts. Instantaneous amplitudes were extracted for all frequencies from 1-30 Hz (1 Hz steps), using a complex Morlet wavelet with a wavelet constant of 8. Afterwards, the 500ms-padding from each side was discarded. For each subject, alpha-band activity was averaged within each condition across 8-12 Hz, across electrodes and across trials, and baseline corrected to a window of 0-100ms post-WS, to avoid contamination by pre-WS systematic differences (see (11)).

To assess the presence of pre-target amplitude suppression, we conducted a t-test against 0, in a pre-defined window of 800-900ms post-WS, where temporal expectation should be maximal. The degree of suppression was compared across conditions through t-tests on planned, directional contrasts with the hypotheses that temporal prediction would lead to stronger suppression, with rhythm leading to the largest effect. We further examined differences in the raw pre-target alpha amplitude with directional t-tests, assessing whether temporal prediction and rhythm in specific would lead to lower pre-target alpha amplitude regardless of degree of suppression.

Lastly, to confirm that the observed suppression was the result of directed temporal attention and not purely stimulus driven, we compared the alpha amplitude in the target interval to that observed in the cue interval, extracted in a matching time window of 800-900ms after the first cue stimulus, and baseline corrected to 0-100ms window after the first cue. This was done for both of the predictive conditions using t-tests for planned contrasts, hypothesizing that the anticipation of the visual target would elicit larger suppression than that of the second cue stimulus.

## Results

To investigate how temporal attention facilitates perception under high perceptual load based on periodic and non-periodic temporal regularities, participants performed a demanding visual discrimination task and reported both the stimulus orientation as well as the subjective visibility. Average discrimination performance across all participants, sessions and conditions was 79%, confirming that the task was perceptually challenging and no ceiling effect was observed.

### Behavior

#### Periodic and aperiodic predictions lead to similar magnitude of perceptual facilitation

To assess the impact of temporal attention on visual perception, psychometric curves were fitted to the aggregated data across sessions of each participant, separately for each condition, and the 75% objective and 50% subjective threshold were extracted (Fig. 2). Repeated measures ANOVAs were conducted separately for subjective and objective measures and revealed a main effect of condition on threshold in both cases (objective: *F*(2,54) = 5.39, *p* = .007, *η²ₚ* = .17; subjective: *F*(2,54) = 7.676, *p* =. 001, *η²ₚ* = .22).

**Figure 2.**
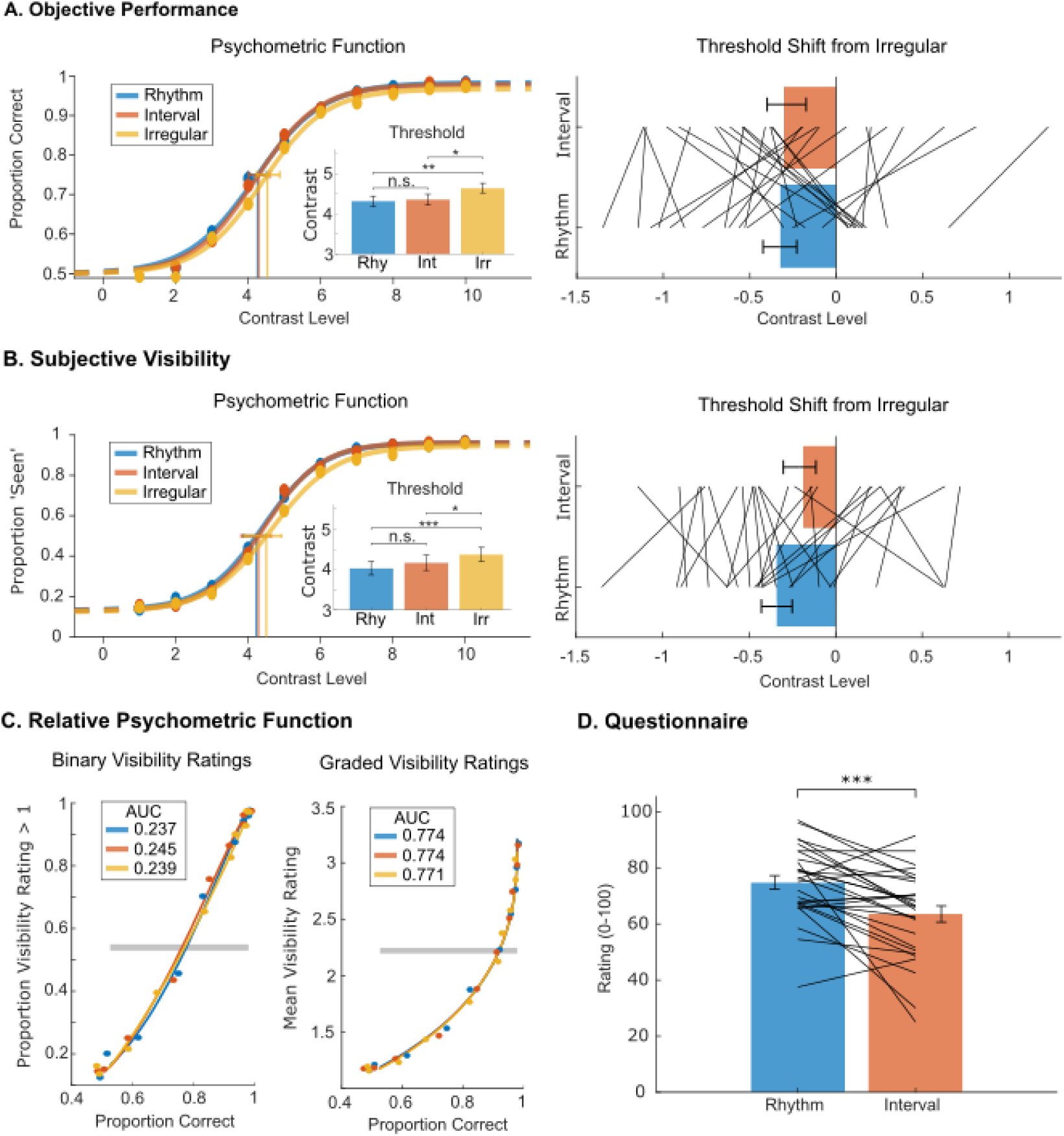
Similar facilitation of objective performance and subjective perception by periodic and aperiodic temporal attention. In all panels: Rhythm (blue), Interval (red), Irregular (yellow). **A.** Psychometric function fits to objective performance. Contrast levels were scaled at participant level. Left panel: Functions are fit to averaged group level performance across target contrast intensities for illustrative purposes. The inlet barplot shows the group averaged mean thresholds (75%) for the three conditions (error bars represent 95% CI). Right panel: threshold shift in the two predictive conditions relative to the irregular condition. Gray lines represent individual participants (error bars represent 95% CI). **B.** Same as A., for subjective performance. To fit the psychometric functions, PAS ratings were binarized (“no experience” rated as 0, otherwise rated as 1). Threshold is at 50% ‘seen’. **C.** Relative psychometric functions (RPF) analysis, with graded (left) and binarized (right) visibility ratings. No effect is found for temporal attention on the relation of objective performance and subjective perception across target contrast intensities. **D.** Post-experiment questionnaire. Responses to the question ‘How well did you manage to use the rhythm (blue)/interval (red)?’. Individual lines representing individual participants, error bars represent 95% CI.

To examine whether timing information improves perception and especially whether rhythmic cues provide an additional benefit, one-tailed t-tests conducted on planned contrasts. In line with previous literature, both temporal predictability significantly lowered the objective perceptual threshold compared to the irregular condition (Fig. 2A; Rhythm: *t*(27) = –3.34, *p* = .001, Cohen’s *d* = –0.63; Interval: *t*(27) = –2.55, *p* = .008, *d* = –0.48). Similar results were obtained for subjective visibility (Fig. 2B; Rhythm: *t*(27) = –3.82, *p* < .001, *d* = –0.72; Interval: *t*(27) = –2.21, *p* = .018, *d* = –0.42). However, no additional benefit of Rhythm was seen compared to Interval in objective performance (*t*(27) = −0.34, *p*= .369, *d* = .06, *BF*=3.8, evidence against stronger effect in Rhythm), although there was a non-significant trend in subjective visibility (*t*(27) = –1.67, *p* = .053, *d* = −.32, one-tailed).

Without a priori hypotheses, we also assessed the effect of timing condition on the slope of the psychometric functions. This was done separately for the objective and the subjective measure, with the slope extracted at the perceptual threshold defined as y=0.75 and y=0.5 respectively. A repeated measures ANOVA revealed no significant slope differences in either the objective (*F*(2,54)=.06, *p*=.935, *η²ₚ* = .003) or the subjective measure (*F*(2,54)=1.19, *p*=.311, *η²ₚ* = .04) between conditions.

#### Relation between Subjective Perception and Objective Performance

We then asked whether and how temporal attention impacts the relation between subjective and objective performance, for example if at specific intensity levels, temporal attention affects one of the measures differently. For this we used the relative psychometric function (RPF; (26)), which allows us to assess the relation between the two measures across the entire psychometric function. Given the slightly different fitting algorithm of the RPF toolbox and consequently different outlier rejections, slight numerical variations in the results compared to the *psignifit* toolbox were expected. However, we confirmed the RPF toolbox replicated the overall results previously obtained with the *psignifit* toolbox for both objective performance (Rhythm-Irregular: *t*(25) = −3.45, *p* (one-tailed) = .001, *d* = −.68; Interval-Irregular: *t*(25) = −2.25, *p (one-tailed)* = .017, *d* = −.44; Rhythm-Interval: *t*(25) = −.86, *p* (one-tailed) = .198, *d* = −.17) and subjective visibility (Rhythm-Irregular: *t*(24) = −2.83, *p* (one-tailed) = .005, *d* = −.57; Interval-Irregular: *t*(24) = −4.29, *p* (one-tailed) < .001, *d* = −.86; Rhythm-Interval: *t*(24) = .25, *p* (one-tailed) = .599, *d* = .05).

Once we confirmed the presence of condition differences for each of the measures, we turned to the key question of the relation between attentional facilitation of objective and subjective perception. A two-tailed repeated measures t-test showed no significant differences on the AUC of the relative psychometric function between any of the conditions (Fig. 2C; Rhythm-Irregular: *t*(22) = .32, *p (two-tailed)* = .753, *d* = .07; Interval-Irregular: *t*(22) = .81, *p (two-tailed)* = .425, *d* = .17; Rhythm-Interval: *t*(22) = −.47, *p (two-tailed)* = .644, *d* = −.10). These results indicate that temporal attention did not alter the relation between the two measures of perception. Repeating this analysis with the graded PAS ratings, as done in a recent study on spatial attention <u>(Tian et al., 2025)</u>, did not result in a meaningfully different pattern for the AUC (Rhythm-Irregular: *t*(25) = .49, *p* (two-tailed) = .626, *d* = .10; Interval-Irregular: *t*(25) = .61, *p* (two-tailed) = .545, *d* = .12; Rhythm-Interval: *t*(25) = −.12, *p* (two-tailed) = .906, *d* = −.02).

#### Post-Experiment Questionnaire

A repeated measures ANOVA on participants’ self-report in the post-experiment questionnaire (“How well did you manage to use the interval/rhythm?”) indicated that following the rhythmic cues was perceived as easier than using the memory-based interval cues (Fig. 2D; *F*(1,28)=17.32, *p*<.001, *η²ₚ*= .72). Additionally, the rating was significantly lower for session 1 than for session 2 (*F*(1,28)=6.84, *p*=.014, *η²ₚ*= .42). There was no interaction between session and timing structure (*F*(1,28)=.04, *p*=.85, *η²ₚ*= .001).

### EEG

#### Delta ITPC

Rhythm-based temporal predictions lead to increased phase alignment of low-frequency neural activity as the predicted target moment approaches in both occipital and frontocentral circuits, typically attributed to oscillatory entrainment to the stream. However, non-rhythmic predictions based on cue-interval associations were shown to lead to similar phase alignment patterns originating in frontocentral circuits in a reaction time task (11). To assess how interval-based predictions impact phase alignment in sensory circuits in the absence of motor demands, we extracted the instantaneous neural phase at the rhythm rate (1.11 Hz), and calculated the inter-trial phase concentration (ITPC) in an occipital electrode cluster. As rhythm entrainment models involve direct impact on sensory circuits, the intuitive hypothesis was that the current task, with its high perceptual but low motor demands, could reveal stronger phase alignment to rhythms specifically in lower sensory regions.

In line with previous results, we observed a significant increase in delta-band ITPC in occipital electrodes during the anticipation interval, clearly diverging 100ms before target onset in the Rhythm condition compared to the Irregular condition (Fig. 3A; *t*(28) = 6.297, *p (one-tailed)* <.001, Cohen’s *d* = 1.169). When comparing the aperiodic Interval condition to the Irregular condition in the same occipital cluster and time window, we also observed increased delta-band ITPC (*t*(28) = 5.984, *p (one-tailed)* <.*001*, *d* = 1.111). Critically, and contrary to the entrainment-based hypothesis, we found no difference in ITPC between the Rhythm and the Interval conditions in the occipital region of interest (*t* (28) = −1.039 *p* = .308, *d* = –.193, *BF* = 9.5, evidence against stronger effect in rhythm), suggesting that non-oscillatory mechanisms can drive sensory phase alignment to an equal degree as rhythmic entrainment.

**Figure 3.**
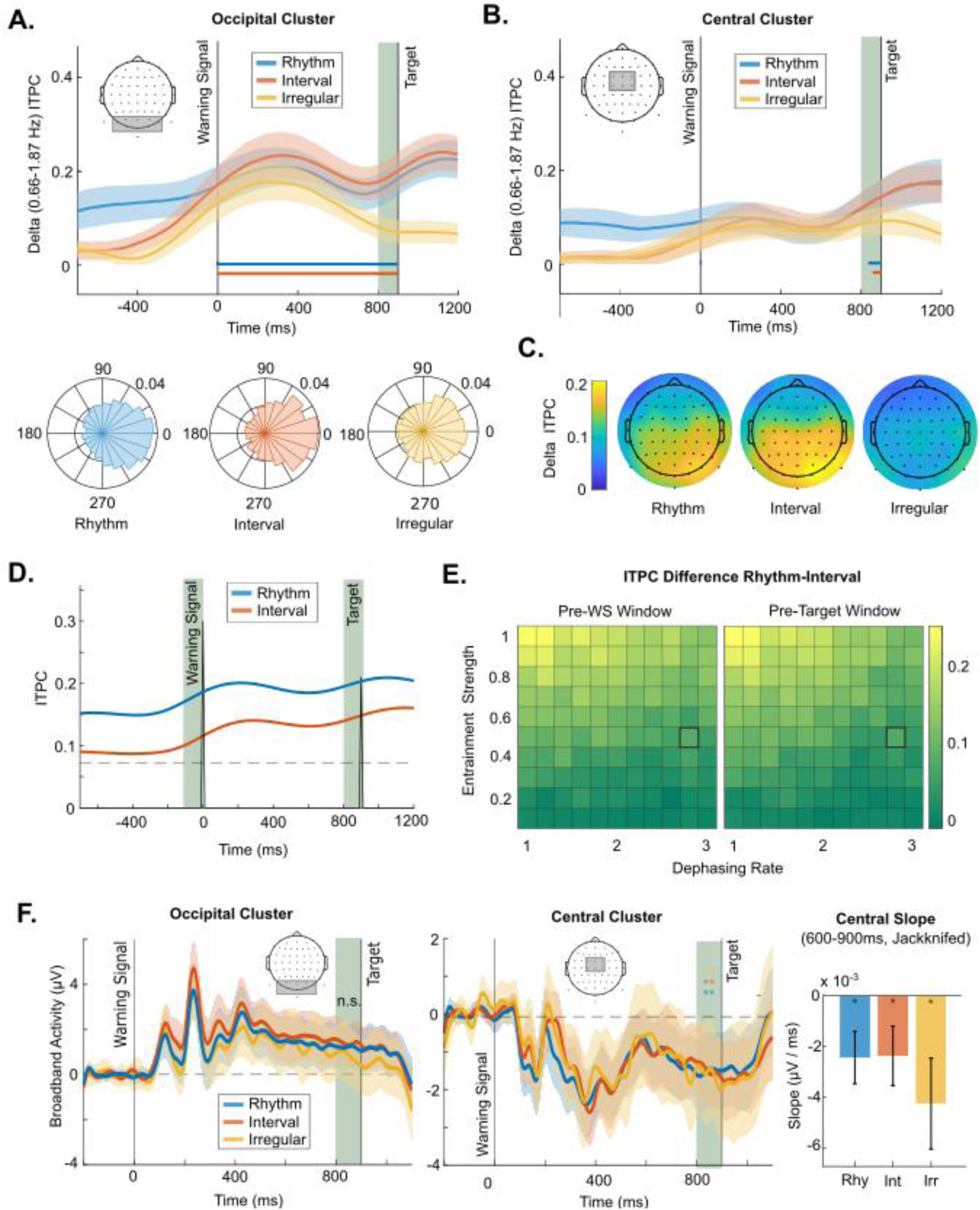
Delta-band (Hz) inter-trial phase concentration (ITPC). **A.** Time-resolved delta-ITPC for the different conditions in occipital and central ROI electrode clusters. Comparable ITPC increase in the predictive conditions in the occipital ROI. **B.** ITPC topography averaged across 800-900ms after the warning signal for each condition, showing an occipital cluster of phase concentration in the Interval and Rhythm conditions. **C.** Phase angle histograms for all trials across participants, separately for each condition. Individual mean is centered on 0 for cross-participant alignment. **D.** Oscillator model simulations of ITPC time course, using parameters fit to the Rhythm condition. An entrainment process predicts weaker pre-target ITPC increase in the Interval condition than in the Rhythm condition, contrary to observed neural data. **E.** ITPC differences between the Rhythm and Interval conditions in the pre-WS and the pre-target window, based on oscillator model simulations across a wide parameter space. No parameter combination leads to ITPC patterns in the observed neural data (large ΔITPC in the pre-WS window, ΔITPC=0 in the pre-target window) based on a purely oscillatory mechanism. **F.** Occipital and central event related potentials in the WS-target window do not agree with ITPC pattern across conditions. Left: no ramping activity in occipital cluster. Middle: ramping activity in a frontocentral cluster, no difference between conditions. Right: Ramping slopes in frontocentral cluster, no difference between conditions (error bars are 95% CI).

Next, we examined delta-band phase alignment in frontocentral circuits, where it has been hypothesized to reflect either anticipatory ramping activity or propagation of the low-level entrainment to motor regions (11,17,34) If frontocentral circuits support a general amodal mechanism of temporal prediction, ITPC should be increased even in the absence of motor preparation. Furthermore, as interval-based predictions do not rely on local dynamics in sensory circuits but on abstract cue-interval associations, we expected stronger involvement of higher-level attentional systems and thus potentially stronger central ITPC in the Interval than the Rhythm condition (34). As in the occipital cluster, we observed higher pre-target ITPC in both predictive conditions compared to the Irregular one (Fig. 3B; Rhythm: *t*(28) = 2.209, *p (one-tailed)=.018*, *d* = .410; Interval: *t*(28) =2.005 *p (one-tailed)* = *.027*, *d* = .372), yet no difference between the two predictive conditions (*t*(28) = −.016, *p (two-tailed)*=.988, *d* = −.003, *BF* = 5.1, evidence in favor of no difference between conditions) in central electrodes. However, inspecting the scalp topographies (Fig. 3C) revealed a single occipital cluster spreading monotonically across the posterior-anterior axis to frontocentral electrodes. Indeed, directly comparing occipital and frontocentral clusters revealed significantly smaller effect of condition (RM ANOVA, Condition X Location interaction, *F*(2,56) = 8.47, *p* < .001) as well as overall weaker ITPC (Location: *F*(1,28) = 4.26, *p* = .048), with planned contrasts showing significantly stronger ITPC in occipital than in central regions in each predictive condition compared to the irregular condition (Rhythm: *F*(1,28) = 8.42, *p* < .001; Interval: *F*(1,28) = 14.34, *p* < .001). These results indicate that the observed central ITPC results from volume conduction from occipital sites, rather than representing a separate activity cluster.

To determine the extent to which the observed ITPC patterns in occipital electrodes, especially in the aperiodic Interval condition, could still be explained by oscillatory entrainment, we simulated the expected oscillatory entrainment with a computational model. We fit the entrainment and dephasing parameters of the model to the observed pre-WS and pre-target ITPC levels in the Rhythm condition, before then using the same parameters to simulate the Interval condition. While we found a parameter combination (dephasing = 2.8, entrainment = 0.5, Fig. 3D) that allowed us to replicate the delta ITPC found in the Rhythm condition and the pre-WS ITPC of the Interval condition, it did not reach the observed pre-target delta ITPC in the Interval condition. This indicates that the observed phase concentration in the Interval condition cannot be explained by oscillatory entrainment to the stimulus stream alone. Further, an exhaustive search of the plausible parameter space (Fig. 3E) revealed that there is no combination which would be able to replicate the observed pattern in the EEG results, of large ΔITPC in the pre-WS window and ΔITPC=0 in the pre-target window. This further supports the notion that top-down factors can drive phase concentration independently of oscillatory entrainment.

Previous work using motor tasks (11,14) has associated an increase in ITPC with broadband anticipatory ramping activity, such as the contingent negative variation (CNV). To assess the relationship between the observed ITPC increase and anticipatory ramping activity, we analyzed event-related potentials in both the occipital and the central regions of interest (Fig. 3F). In the occipital cluster, the broadband amplitude in the 100ms pre-target window showed no anticipatory negative ramping, and was in fact significantly more positive than baseline (Rhythm: t(28) = 2.563, p = .016, *d* = .476; Interval: t(28) = 2.995, p = .006, *d* = .556; Irregular: t(28) = 1.340, p = .191, *d* = .249), reflecting carryover from the positive evoked response (compare with (11)). Instead, in central electrodes, the pre-target amplitude was more negative in all conditions compared to baseline (Rhythm: t(28) = −3.261, p(one-tailed) = .002, *d* = .606; Interval: t(28) = −3.363, p(one-tailed) = .001, *d* = .625; Irregular: t(28) = −2.658, p(one-tailed) = .006, *d* = 0.494), with no difference between conditions (Rhythm-Irregular: t(28) = .823, p = .417, *d* = .153; Interval-Irregular: t(28) = 1.722, p = .096, *d* = .320; Rhythm-Interval: t(28) = −.910, p = .370, *d* = −.169). Moreover, there was a significant negative slope in the 300ms before target onset in all conditions (Rhythm: t(28) = −2.375, p = .012; Interval: t(28) = −2.031, p = .026; Irregular: t(28) = −2.276, p = .012). These findings are inconsistent with the selective ITPC enhancement observed in the predictive conditions, indicating a dissociation between anticipatory ramping activity and delta ITPC.

#### Alpha Amplitude

Both spontaneous fluctuations and directed changes in occipital alpha power have been associated with changes in visual detection, with lower alpha amplitude being linked to facilitated visual processing (35–37). Meanwhile, anticipation of an incoming visual target has been consistently shown to lead to targeted suppression of occipital alpha power in spatial attention paradigms (16,38,39) as well as both periodic and aperiodic temporal attention studies (15,40,41) providing a potentially mechanism through with attention modulates visual perception.

To assess whether our data show anticipatory alpha suppression, we first compared alpha amplitude in a 100ms-window pre-target to a 100ms-window post-warning signal within each condition (Fig. 4A). We observed strong pre-target alpha suppression in all conditions (Rhythm: *t*(28) = −5.013, *p (one-tailed)<.001*, Cohen’s *d* = –0.931; Interval: *t*(28) = −4.398, *p (one-tailed) <.001*, *d* = −.817; irregular: *t*(28) =-5.091, *p (one-tailed) <.001*, *d* =-.945). Strikingly, and inconsistent with the behavioral modulations, temporal prediction did not lead to a larger magnitude of suppression at the pre-target window relative to the Irregular condition (Fig. 4B; *t*(28) = −1.048, *p (one-tailed) = .152*, *d* = *−.195*, *BF =* 1.9), with Interval even being slightly less negative than irregular (*t*(28) = .896, *p = 0.378*, *d* = .166, *BF* = 8.8, evidence in favor of no suppression relative to Irregular). Even though there was a trend towards stronger alpha suppression in Rhythm compared to Interval (*t*(28) = −1.858, *p =.073*, *d* = *−.345*), this cannot be interpreted as temporally predictive effect given the lack of difference from irregular (compare with previous work (15,42). Consistent with these results, a cluster-based permutation analyses (10000 permutations) found no clusters of significant difference at any time point between conditions.

**Figure 4.**
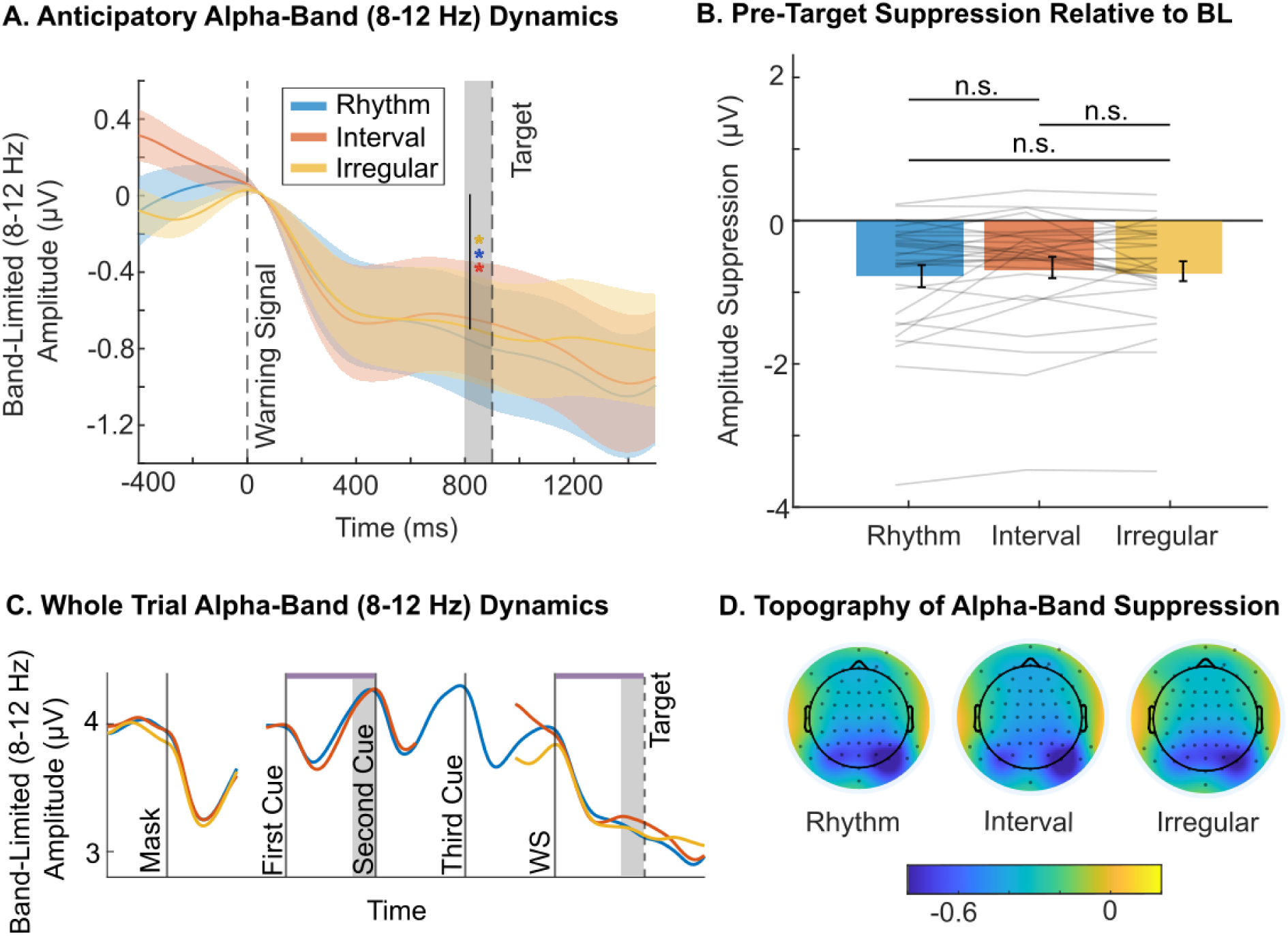
Anticipatory suppression of alpha-band (8-12 Hz) amplitude. **A.** Alpha-band amplitude dynamics in the WS-target window, for the three experimental conditions, baseline-corrected to 0-100ms post-WS. Anticipatory suppression is significant in all conditions (* p<0.001), whereas there is no difference in the degree of suppression between conditions. **B.** Pre-target suppression relative to baseline (0-100ms post-WS) for each condition; Rhythm (blue), Interval (orange) and Irregular (yellow). Grey lines represent individual participants, error bars represent 95% CI. **C.** Whole trial dynamics of alpha amplitude (without any baseline correction) for the three experimental conditions Rhythm (blue), Interval (orange) and Irregular (yellow). Temporal attention in the pre-target interval leads to stronger amplitude suppression than the stimulus-evoked amplitude suppression in the first-second cue interval. **D.** Topography of alpha-band amplitude compared to baseline, averaged across 800-900ms post WS. Alpha suppression is primarily occurring in occipital electrodes across conditions.

One possible explanation to this lack of difference between the conditions is that even under the high perceptual demands of the paradigm, the observed suppression does not reflect proactive increase of attention but purely stimulus-driven suppression to the presentation of the WS. To reject this possibility, we examined alpha-band amplitude dynamics across the whole trial (Fig. 4C), and compared alpha amplitude dynamics between the WS-target interval and the first cue interval, in which the timing of the upcoming stimulus was predictable but not requiring proactive temporal attention. Indeed, after some initial suppression alpha amplitude rebounds to above baseline, leading to significantly stronger suppression in anticipation of the target than the second cue-stimulus (Rhythm: *t*(28) = −4.916, *p (one-tailed) < .001*, *d* = *−.91;* Interval: *t*(28) = −4.152, *p (one-tailed) < .001*, *d* = −.77). Thus, the alpha suppression observed following the WS cannot be explained by a passive stimulus response, and is more likely to reflect guided temporal attention to target onset.

Further, though we chose a 100ms post-WS window as a baseline, to avoid contamination by systematic pre-WS condition differences (Breska et al. 2017, see methods), we repeated our analysis on data which was baseline-corrected to a 100ms pre-WS to ensure that potential differences between conditions were not obscured by the choice of baseline. Indeed, with neither baseline did we observe a difference in pre-target alpha amplitude between conditions.

## Discussion

Temporal predictions that allow for targeted shifting of attention in time can be generated through various periodic and aperiodic temporal regularities in the environment. In the current study, we directly compared how temporal predictions generated through either regularity impact visual perception and anticipatory neural dynamics in visual circuits, while substantially reducing interference of motor preparation. We found that relative to a non-predictive irregular stream, aperiodic cue-interval-based predictions led to similar perceptual facilitation and, strikingly, to similar increase of delta-band phase alignment as rhythm-based predictions, an effect that could not be explained by a time-resolved computational entrainment model. In contrast, anticipatory suppression of alpha-band amplitude was similar in all conditions, diverging from the behavioral results. Altogether, these results uncover a unique spectral profile of temporal visual attention, highlight phase control as a key mechanism, and reveal the potency of discrete cue-interval regularities in driving sensory phase alignment.

### Temporal attention improves perception across measures and temporal structures

The core rationale of our study was that periodic and aperiodic regularities might differ in how they affect pure perceptual facilitation. In periodic streams, oscillatory entrainment occurs in sensory circuits themselves, driving prediction through limit cycle dynamics (3,43). In contrast, in aperiodic cue-interval regularities, the expected interval is not present in local sensory dynamics. Instead, prediction depends on the coordination of a broad network of timing, memory, and attentional circuits, potentially leading to a noisier prediction process and weaker predictive facilitation (44–47). While previous studies found similar behavioral facilitation and neural phase alignment for rhythm- and interval-based predictions (11) this could result from the use of a reaction time task, which could have obscured the sensory-specific advantage of rhythms and lead to alignment of well-described ramping activity in motor circuits (12,14,32).

To maximize sensitivity to behavioral expressions of this difference and obtain a more fine-grained look into the effects of temporal attention on perception, we measured both objective and subjective perception, across a broad range of target contrasts to probe the entirety of the psychometric function. Consistent with prior studies (e.g. (2,48)), we observed a significant reduction of the objective discrimination threshold and of the subjective perception threshold for both predictable conditions compared to the Irregular one. Crucially, both interval and rhythm cues lowered each of the objective and subjective thresholds to a comparable degree. Thus, in contrast to the entrainment benefit perspective, even in a perceptually demanding task there is no additional performance benefit of rhythmic entrainment above and beyond isolated temporal expectation.

Notably, we found no effect of temporal attention on the slope of the psychometric function in either regularity type for either measure (for similar findings for objective performance in rhythm-based prediction, see (4)). The lack of slope effect in combination with a shift in objective threshold was suggested to reflect temporal attention affecting perceptual performance through contrast gain, rather than response gain (49,50). A threshold shift without a change in slope for subjective visibility could be the result of either an improved signal-to-noise ratio, or of an increased bias to report vague stimuli as ‘seen’. While our paradigm does not allow us to clearly distinguish between these two possibilities, the results of the relative psychometric function suggest that temporal attention does not alter the relation between objective and subjective performance, which is consistent with enhancement of sensory evidence, rather than bias.

The overall similar facilitation by temporal attention of both objective performance and subjective visibility across both cue types stands in contrast to findings from other types of attention. Specifically, several spatial attention or object-based attention studies reported a dissociation between the attentional impact on objective performance and subjective visibility or confidence (spatial: (51,52)) object-based: (53)). Additionally, a recent study (27) showed that the relationship between objective perception and subjective visibility is altered under spatial attention, which we did not find for temporal attention. These discrepancies suggest that temporal attention might affect perception in a fundamentally different manner than other types of attention.

Interestingly, in our post-study questionnaire, participants reported that following the rhythmic cues was perceived as easier than following the interval cues. The distinction between the similar perceptual benefits of both conditions and the lower perceived subjective effort for rhythm-based prediction might indicate that interval timing can reach the performance benefits of rhythmic timing only when substantial mental effort is employed. Future research could test this hypothesis by investigating whether memory-based predictions fail to match rhythmic performance as mental fatigue increases or when working memory is occupied, as working memory load has been shown to impact interval timing (54,55).

### Aperiodic Interval predictions lead to increased occipital delta ITPC

Our findings of similar behavioral facilitation by rhythm- and interval-based predictions do not necessarily imply reliance on overlapping neural mechanisms. For example, previous work showed that although they lead to similar response time facilitation, they are supported by distinct subcortical circuits (56,57). Following the same core rationale as above, we hypothesized that rhythm-based prediction would lead to stronger occipital delta-band phase alignment through entrainment of the sensory cortex than interval-based prediction. However, in occipital electrodes, we found that interval-based predictions increased ITPC stronger than the irregular condition and, surprisingly, also that the magnitude of the ITPC increase was similar to that driven by rhythm-based prediction.

Notably, one previous study reported increased ITPC in an interval-based explicit timing task (earlier/later than expected) compared to luminance judgments, attributing it to phase-resetting of ongoing delta-band oscillations (31). Here we test the impact of interval-based prediction against an irregular condition, in which time is still relevant but with higher uncertainty that does not allow for precise temporal prediction. In line with this study, our time-resolved oscillatory entrainment model does indicate that phase reset to the onset of the predicted interval is sufficient to drive some ITPC increase. However, our simulations show that this phenomenon alone should not match the magnitude of the ITPC increase by isochronous streams. Thus, in a perceptually demanding context, memory-based predictions can lead to significant top-down phase alignment even in lower sensory cortices, optimally preparing the system for perceptual demands in the absence of entrainment.

Delta-band phase alignment by temporal predictions has also been observed in fronto-central circuits, but the use of motor tasks precluded from attributing them to top-down control of sensory processing or to motor preparation (11). Indeed, previous work using visual rhythms found no evidence of phase alignment in central sites (48). Our design allowed us to test the hypothesis of stronger fronto-central delta phase alignment in the Interval condition than the Rhythm condition, as the former putatively relies more on top-down attentional control.

However, in contrast to this hypothesis, we found no significant increase of delta ITPC in central electrodes in any of the conditions. These results suggest that the previously observed delta-band ITPC in central sites in a visual speeded responses task reflects motor processes rather than pure timing mechanisms. More broadly, they suggest that top-down temporal attention in interval-based prediction and its phase alignment outcomes in sensory circuits is not supported by phase alignment in corresponding frequencies in central regions. Furthermore, these results are not in line with the idea that the motor system is supporting rhythm-based sensory prediction, though this connection is hypothesized to be more fundamental in auditory rhythms (17).

Our modelling results show that phase alignment in the Interval condition cannot be explained by oscillatory entrainment alone, raising the question of their origin. One previously suggested hypothesis is that delta phase alignment can reflect anticipatory ramping activity (11). Of note, this was suggested for central delta phase alignment in speeded response tasks, with ramping activity much more prevalent in frontal and motor circuits than in sensory circuits (though (58). However, our ERP analysis revealed that the increased delta ITPC in occipital electrodes was not accompanied by clear ramping activity in the broadband response.

Surprisingly, even in the absence of low-frequency ITPC, we do find neural activity in central electrodes that appears to show the typical negative ramping in anticipation of the target (12,14,32), before resolving after the moment of target presentation. Notably, as the current paradigm did not include variable ISIs between warning signal and target, it is difficult to determine whether the observed central ramping activity is flexibility adjustable with goal-directed attention, or rather merely a passive byproduct of stimulus evoked responses propagated forward. Nevertheless, these overall results suggest that the observed delta ITPC is unlikely to be a necessary outcome of consistent anticipatory ramping activity. We suggest that it might instead represent a separate aperiodic mechanism of top-down temporal attention. Furthermore, they corroborate previous suggestions that low-frequency phase concentration is not a unique marker of rhythmic entrainment (11,30).

### Behavioral benefit of temporal attention is not reflected in alpha-band suppression

Posterior alpha-band power has been related to visual perception (35–37) and is thought to reflect a mechanism of attention through inhibition (59). Several studies of interval-(40,41) or rhythm-based predictions (15) showed anticipatory suppression in alpha power prior to predicted target onset, interpreted accordingly as reflecting the optimization of the visual system for the perception of the upcoming stimulus at the expected time. We tested the hypothesis that compared to rhythm-based prediction, interval-based prediction might be associated with stronger alpha suppression to compensate for the lack of local entrained dynamics, akin to its known role in endogenous, cue-based shifting of spatial attention (16,38,39). Inconsistent with this hypothesis, while we did find significant suppression of alpha power prior to target onset relative to baseline, we found similar magnitude of suppression in the Interval and Rhythm conditions. Considering that interval-based prediction led, in our paradigm, to similar delta-band alignment, such compensation is perhaps not necessary.

However, this interpretation should be made cautiously as the strength of the suppression did not vary with temporal predictability of the target. Instead, we see a significant suppression also in the Irregular condition, in which the onset of the target stimulus had low predictability.

This result stands in contrast to previous findings of greater anticipatory alpha suppression compared to a non-predictive condition (15,40,41). Further, as far as it was possible to reconstruct alpha-band activity across the whole trial, we found no significant difference across conditions in the whole trial dynamics. A series of control analyses rejected potential alternative confounding explanations. First, the WS-target ISI was not too short to allow for recovery of the signal to baseline in the Irregular condition, as stimulus-evoked suppression of the cue stimuli did return to baseline as seen in the activity across the whole trial. Second, the observed suppression in the WS-target interval was not solely stimulus driven, as it was stronger than in the first cue interval of the predictive conditions, even though in the latter the timing of the second stimulus was as predictable. Thus, it appears that alpha activity could be equally suppressed in active anticipation of the target in all conditions, even in the absence of precise temporal information.

Speculatively, one explanation for this result is that, contrary to delta phase alignment, alpha suppression could represent a less precise timing mechanism. Thus, the warning signal in the irregular condition might have provided enough temporal information for anticipatory alpha suppression to occur. This idea is supported by the dynamics of the suppression being temporally broad, rather than precisely timed to the expected moment. The discrepancy from previous work finding alpha suppression in temporal prediction could stem from using a less demanding visual task might have not driven strong need for suppression (but see (40)), or from combining spatial-temporal cuing which could have further engaged alpha-based spatial-related mechanisms.

Moreover, previous studies often compared alpha levels at an expected time to expected an event at a different time point, potentially reflecting active inhibition of attention (11,15,60). Our findings suggest that under high visual load and no spatial uncertainty, the magnitude of temporally cued alpha-band suppression is not stronger than a more ‘neutral’ state of distributed attention in time.

Importantly, regardless of the explanation for the similar degree of suppression in the predictive and non-predictive conditions, this pattern is discrepant from the significant behavioral benefits we observed in both objective performance and subjective visibility. Thus, the current findings suggest that in contrast to the traditional view (61), alpha suppression might not be the mechanism responsible for the behavioral effects observed, pointing to the distinct neural organization of temporal attention. Regarding more recent studies suggesting that alpha amplitude is mainly related to shifts in response criterion rather than visual sensitivity ((62–64) but see (65)), it is important to note that in the current study we observe a shift in subjective visibility under temporal attention in the absence of condition-specific alpha modulation. This further supports our prior speculation that the threshold shift we observe in subjective visibility is likely due to improved perception, rather than a shift in response criterion. It should be noted that our design might have been suboptimal to investigate alpha activity due to the continuous dynamic noise mask, as well as the difference in luminance of cue and warning stimuli.

### Summary

Overall, our results show that even in a challenging perceptual task, top-down aperiodic temporal predictions can drive both behavioral benefits and neural responses to an equal degree as entrainment-based temporal attention. In contrast to consistently reported subcortical dissociations, the current findings call into question whether different temporal structures rely on different neural mechanisms, at least in cortical sensory circuits. We also show that temporal attention can lead to an increase in low-frequency phase alignment in sensory regions even in the absence of neural entrainment. Conversely, phase alignment in central regions reported in prior study does not appear to be a necessary component of temporal attention and associated ramping activity. Furthermore, the current results dissociate anticipatory alpha suppression from delta phase alignment as well as the perceptual benefits generated by temporal attention.

## Acknowledgements

This research was supported by funding from the Max Planck Society (Max Planck Independent Research Group) to Assaf Breska.

